# A low-dimensional approximation of optimal confidence

**DOI:** 10.1101/2023.03.15.532729

**Authors:** Pierre Le Denmat, Tom Verguts, Kobe Desender

## Abstract

Human decision making is accompanied by a sense of confidence. According to Bayesian decision theory, confidence reflects the learned probability of making a correct response, given available data (e.g., accumulated stimulus evidence and response time). Although optimal, independently learning these probabilities for all possible combinations of data is computationally intractable. Here, we describe a novel model of confidence implementing a low-dimensional approximation of this optimal yet intractable solution. Using a low number of free parameters, this model allows efficient estimation of confidence, while at the same time accounting for idiosyncrasies, different kinds of biases and deviation from the optimal probability correct. Our model dissociates confidence biases resulting from individuals’ estimate of the reliability of evidence (captured by parameter α), from confidence biases resulting from general stimulus-independent under- and overconfidence (captured by parameter β). We provide empirical evidence that this model accurately fits both choice data (accuracy, response time) and trial-by-trial confidence ratings simultaneously. Finally, we test and empirically validate two novel predictions of the model, namely that 1) changes in confidence can be independent of performance and 2) selectively manipulating each parameter of our model leads to distinct patterns of confidence judgments. As the first tractable and flexible account of the computation of confidence, our model provides concrete tools to construct computationally more plausible models, and offers a clear framework to interpret and further resolve different forms of confidence biases.

**Significance statement:** Mathematical and computational work has shown that in order to optimize decision making, humans and other adaptive agents must compute confidence in their perception and actions. Currently, it remains unknown how this confidence is computed. We demonstrate how humans can approximate confidence in a tractable manner. Our computational model makes novel predictions about when confidence will be biased (e.g., over- or underconfidence due to selective environmental feedback). We empirically tested these predictions in a novel experimental paradigm, by providing continuous model-based feedback. We observed that different feedback manipulations elicited distinct patterns of confidence judgments, in ways predicted by the model. Overall, we offer a framework to both interpret optimal confidence and resolve confidence biases that characterize several psychiatric disorders.

## Introduction

Decision confidence refers to a subjective feeling reflecting how confident agents feel about the accuracy of their decisions. This feeling of confidence closely tracks the objective accuracy (1): people usually report high confidence for correct trials and low confidence for errors. This observation is in line with the theoretical proposal that confidence reflects the Bayesian posterior probability that a decision is correct given available data (1–3). As such, confidence represents valuable information that is taken into account to guide adaptive behavior, including learning (4–6); speed-accuracy tradeoff adjustments (7, 8); and information seeking (9). Therefore, having an accurate sense of confidence that best matches one’s accuracy is of utmost importance to maintain adaptive behavior. Dissociations between confidence and accuracy are widespread, however, most prominently in cases of blindsight (10), change blindness (11) and patients with anterior prefrontal lesions (12). Such dissociations pose a serious challenge for the Bayesian interpretation of confidence. More importantly, estimating the Bayesian probability with limited data is computationally intractable. In this work, we reconcile these findings by proposing and empirically validating a low-dimensional approximation to the Bayesian probability, offering both a tractable and flexible model for the computation of decision confidence.

Most attempts at modeling decision confidence have done so within the context of existing models of decision making. One highly influential account is based on the idea that decision making reflects a process of noisy accumulation of evidence until a decision boundary is reached (13). For example, the drift-diffusion model (DDM) describes the decision-making process as the noisy accumulation of evidence in favor of one of two options. Here, evidence accumulates with a certain drift rate (representing the efficiency of evidence accumulation) until reaching a decision threshold, at which point a response is issued. Several approaches have been put forward to account for confidence within the DDM framework (14–16). The most prominent approach relies on the Bayesian interpretation of confidence, modeling it as the probability of a choice being correct given the available data. Within the drift diffusion model, the available data to participants is the amount of accumulated evidence and the time spent accumulating, which are then combined into a probability that the decision was correct (2, 15). Such formalization of decision confidence is sometimes referred to as the “Bayesian readout” (17). This Bayesian readout can be represented as a heatmap on the two-dimensional (data) space formed by both evidence and time. In Figure 1A, it can be seen that the Bayesian readout hypothesis predicts that confidence will be higher for trials with more accumulated evidence (reflected on the y-axis) and lower for trials with a longer decision duration (reflected on the x-axis). Consistent with these predictions, confidence indeed depends on evidence strength (1, 2) and on elapsed decision time (14). More generally, this modeling approach has been successful in explaining a wealth of data (17–19).

**Figure 1.**
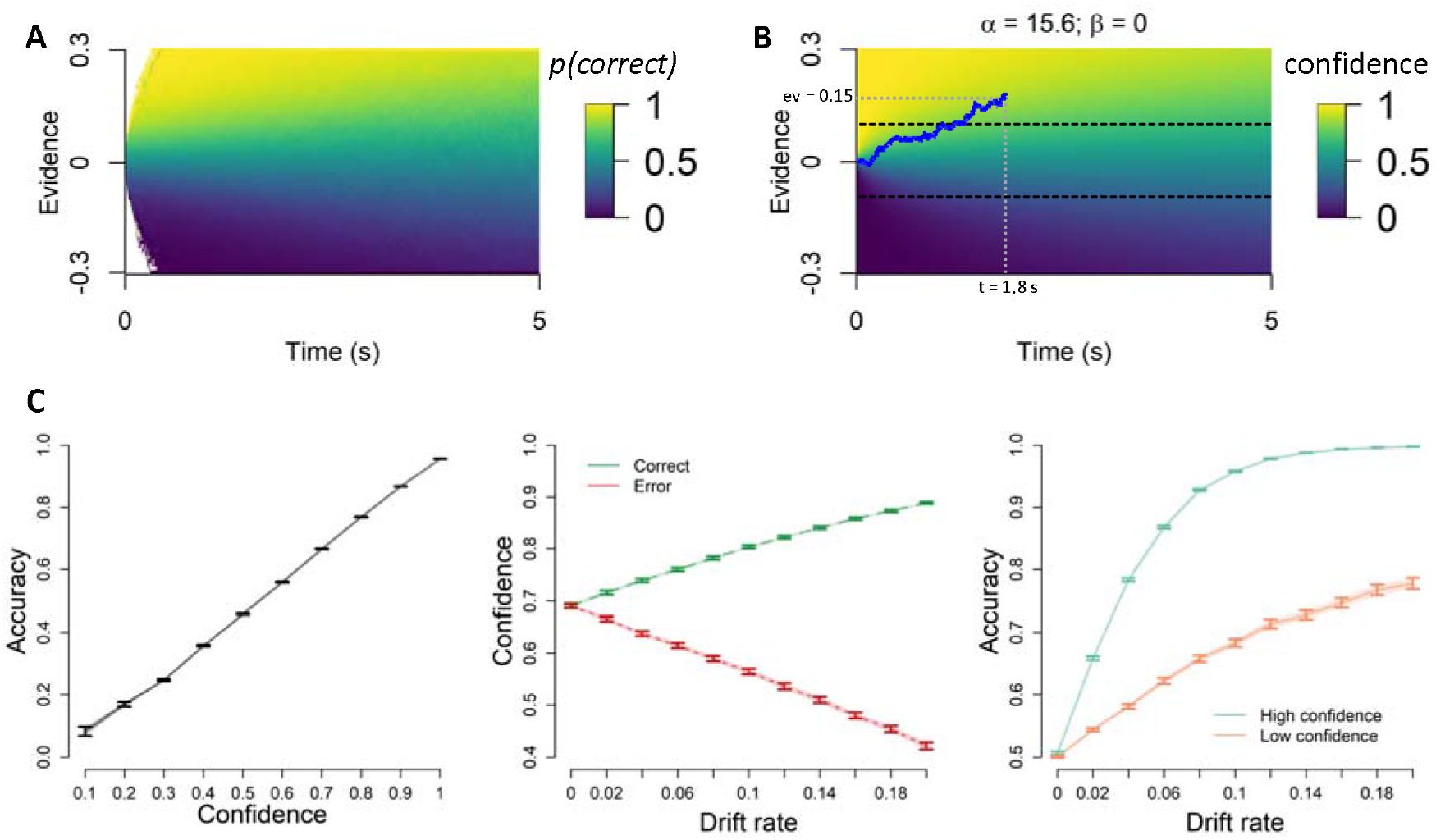
**A.**Confidence is thought to represent the Bayesian probability of a choice being correct conditional on evidence, time and choice. Within this theory, confidence is quantified as this probability, represented by the color on the heatmap. **B**. Because this optimal solution is intractable, the LDC model proposes a low-dimensional parametrization of this framework, which allows efficient estimation of confidence, while accounting for idiosyncrasies and confidence biases. The LDC model can generate a heatmap representing confidence which closely approximates the optimal Bayesian probability. Values of α and β were obtained by fitting the LDC model to the Bayesian probability of being correct over 1 000 000 simulated trials. Confidence for the trial plotted on top of the heatmap is given by Eq. (3). Here, confidence = .85. **C**. To show the effectiveness of the LDC model we generated three statistical signatures of confidence (Sanders et al. 2016) based on the Bayesian read-out of confidence (error bars reflecting SEM, simulated N = 100) and based on the LDC model fits (shaded lines reflecting SEM).

To compute confidence by reading out the probability correct given evidence and time, humans must have an accurate representation of the entire space created by crossing these two variables (i.e. the heatmap shown in Figure 1A). Previous accounts propose that individuals learn this mapping via experience (2). However, learning all positions on this heatmap would either take a lot of time or yield very noisy estimates. Thus, tractability is a key issue that needs to be addressed in order to understand how humans learn the probability correct given evidence and time. Therefore, in the current work we propose the Low-Dimensional Confidence (LDC) model, a simple yet efficient low-dimensional approximation of the optimal yet intractable Bayesian readout. In the following sections, we describe how LDC allows to tractably compute the mapping from evidence and time to confidence. Using simulated data, we show that LDC provides a close approximation of Bayesian confidence. We then proceed to test and validate our model with human data.

## Results

### The Low-Dimensional approximation of Confidence model (LDC model)

Constructing an accurate representation of confidence based on a limited number of samples is infeasible. However, under standard DDM assumptions, the probability of a correct choice given accumulated evidence and elapsed time can be expressed as the probability of drift rate v being positive in case of upper boundary hit (and conversely *p*(*ν* < 0) in the lower boundary hit case). Such probability is characterized as (15):

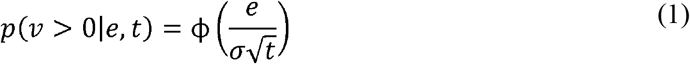

where *e* is the accumulated evidence, *t* is the elapsed time, ϕ is the cumulative distribution function of the standard normal distribution and *σ* is the within-trial noise of the DDM accumulator Given that ϕ is an integral without closed-form solution, it requires an infinite number of standard operations to be computed. We propose to approximate ϕ with a more tractable logistic function (20):

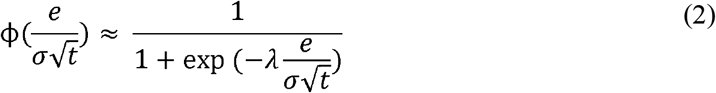

where *λ* ≈ 1.7 is a constant that optimizes the approximation (20). In its current form, the formalization of confidence proposed in Eq. (2) cannot account for idiosyncrasies (21), diverse types of confidence biases and deviations from the optimal probability of a correct choice typically observed in empirical work (22–24). In order to make the formulation of confidence more flexible we thus further parameterize confidence in the following way:

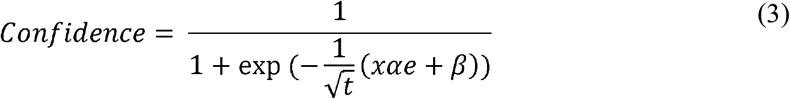

where *x* ∈ {−1; 1} is the choice. The two free parameters of this equation capture how strongly individuals weigh evidence in their computation of confidence (α); and a stimulus-independent confidence bias (β). A positive confidence bias (β > 0) implies that the model has a general tendency to be overconfident. If β = 0, the model is unbiased and bases its confidence purely on the evidence accumulated and the time spent accumulating. A negative confidence bias (β < 0) indicates overall underconfidence.

As a weighting parameter on evidence, α can be interpreted as individuals’ estimate of the reliability of evidence. Intuitively, if one thinks that the accumulated evidence is not reliable, one will need more evidence to be sure that the decision was correct. When α is decreased, one puts less importance on accumulated evidence to compute confidence. In the extreme case where α = 0, the model completely ignores evidence and the computation of confidence is entirely driven by β and time. If additionally β = 0, then confidence will always be .5. At the other end of the spectrum, if α tends to infinity, then the smallest amount of evidence will lead to extreme confidence judgments (i.e. either confidence = 1 if *ev* > 0 or confidence = 0 if *ev* < 0). Given that accumulated evidence is noisy, an individual with an overly high α likely treats evidence as more reliable than it actually is.

### Simulations: The LDC model closely resembles Bayesian confidence

The aim of the current work is to provide a tractable and flexible approximation of the Bayesian readout of confidence. A first test of the LDC model is whether it can effectively approximate the Bayesian readout of confidence. For this sake, we generated data from 100 simulated participants from a range of typically observed DDM parameters. Our model was then fit to the true Bayesian posterior probability correct conditional on evidence, time and choice. LDC- predicted confidence almost perfectly correlated with the true probability of being correct (Spearman r(999998) = .99, p < .001). This close resemblance can be appreciated visually by comparing the model-based heatmap (created based on the estimated parameters; Figure 1B) to the heatmap based on the simulations (Figure 1A).

To further show that our model closely tracks the Bayesian readout of confidence, we tested its ability to reproduce three statistical signatures that confidence should adhere to if it does reflect a Bayesian probability (Sanders et al., 2016). The three qualitative signatures are 1) confidence predicts choice accuracy, 2) confidence increases with evidence strength for correct trials, but decreases with evidence strength for error trials (commonly called the folded X-pattern; 25, 26) and 3) for any level of evidence strength above 0, high confidence trials should be linked with higher accuracy than low confidence trials. As can be assessed on Figure 1C, the simulated data showed an excellent fit to the signatures.

### Empirically testing predictions of the LDC model

Having demonstrated that the LDC model can closely approximate the Bayesian readout of confidence on synthetic data, we next turned to empirical data from human participants. We tested two key predictions of the LDC model. First, the LCD model predicts that changes in confidence can be independent of performance. The two free parameters only describe how evidence and time are combined into a confidence judgment, but they do not affect the process that leads to specific levels of accumulated evidence and elapsed time. Any manipulation that selectively targets confidence while leaving performance unaffected should thus be captured by changes in α and/or β. A second novel prediction is that selective changes in each parameter of our model should lead to distinct modulations of confidence judgments. Thus, a manipulation targeting reliability (α) should lead to qualitatively distinct changes in confidence ratings compared to a manipulation targeting confidence bias (β).

#### Experiment 1: The LDC model accounts for performance-independent changes in confidence

We first tested a crucial prediction of our model, namely that changes in confidence can occur independent of changes in performance (9, 27–29). Although such dissociations have been observed since several decades (e.g., blindsight; Weiskrantz et al., 1974), they pose a serious challenge for most current models of confidence. The LDC model naturally accounts for such dissociations. One particularly strong dissociation was observed in our recent work (19), in which a manipulation of participants’ prior belief about their ability to perform a task selectively influenced their reported levels of confidence. In Experiment 1 of that paper, participants performed three perceptual tasks consecutively, each divided into a training and a testing phase (Figure 2). During the training phase, participants received feedback about their performance every 24 trials. Although participants were told that the feedback indicated how well they performed the task compared to a reference group, in reality the feedback was made up. Within each task, feedback indicated that performance was worse than of most other participants *(negative condition);* that it was on average *(average condition);* or that it was better than of most other participants *(positive condition)*. During the testing phase, participants no longer received feedback; instead, they rated their confidence at the end of each trial. We observed a direct influence of the feedback manipulation on confidence, with more positive feedback leading to higher confidence, *F*(2,47) = 16.65, *p* < .001. Importantly, this effect of feedback on confidence was not explained by objective performance, as reaction time (RT) and accuracy did not change as a function of feedback (accuracy: *X*^2^(2) = 0.3, *p* = .863; RT: *F*(2,48) = 2.06, *p* = .14).

**Figure 2.**
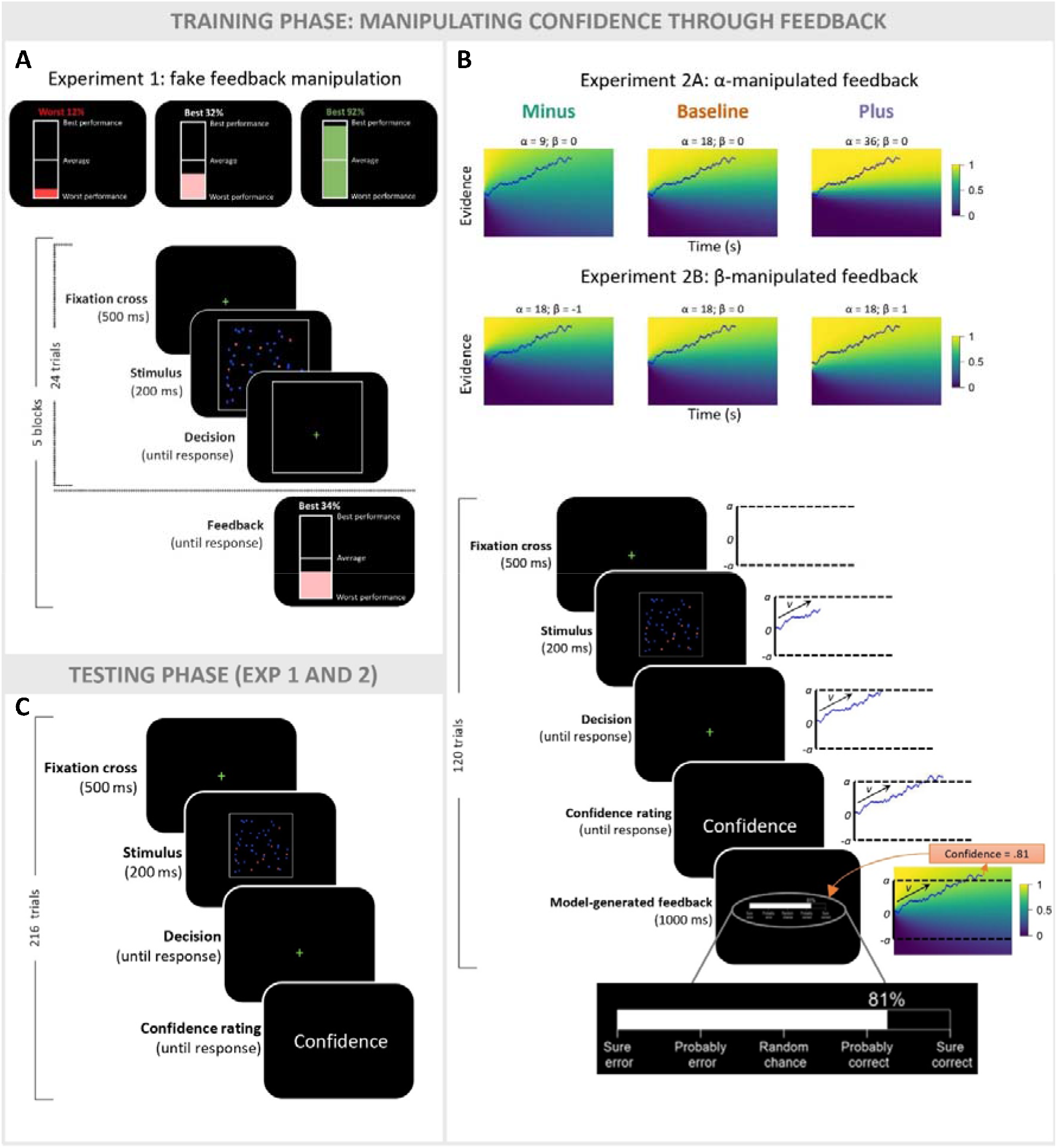
Experimental design. In both experiments, participants performed three different perceptual decision-making tasks (only one shown here). Each task started with a training phase during which a different feedback manipulation was induced. **A**. In Experiment 1, participants received fake feedback after each training block, framed as a comparison between their performance and the performance of a reference group. **B**. In Experiment 2, participants additionally rated their confidence before receiving trial-by-trial feedback reflecting their probability of making the correct choice. Unknown to participants, the feedback was actually generated by the LDC model behind the curtain. To do so, the evidence accumulation process for each trial was estimated using the mean drift rate and boundary from a previous pilot session (see Methods for full details). Feedback conditions differed in the α (resp. β) value used to generate feedback in Experiment 2A (resp. Experiment 2B). **C.**In both experiments, after each training phase participants completed a test phase during which they no longer received feedback but rated their decision confidence after each decision.

We fitted the LDC model to the performance (accuracy and RT) and confidence reports in the test phase of this experiment, separately for each participant. LDC model predictions were then generated using the best fitting parameters for each individual. As can be seen in Figure 3, the LDC model provided an excellent fit to the data. Similar to the empirical data, feedback significantly influenced model-generated confidence ratings (F(2,48) = 9.79, p < .001), but did not influence the performance data (RT: F(2,48) = 1.19, p = .31; Accuracy: *X*^2^(2) = .75, p = .69). Thus, our model was able to capture the data pattern, namely that confidence reports can be influenced independently from behavioral performance.

**Figure 3.**
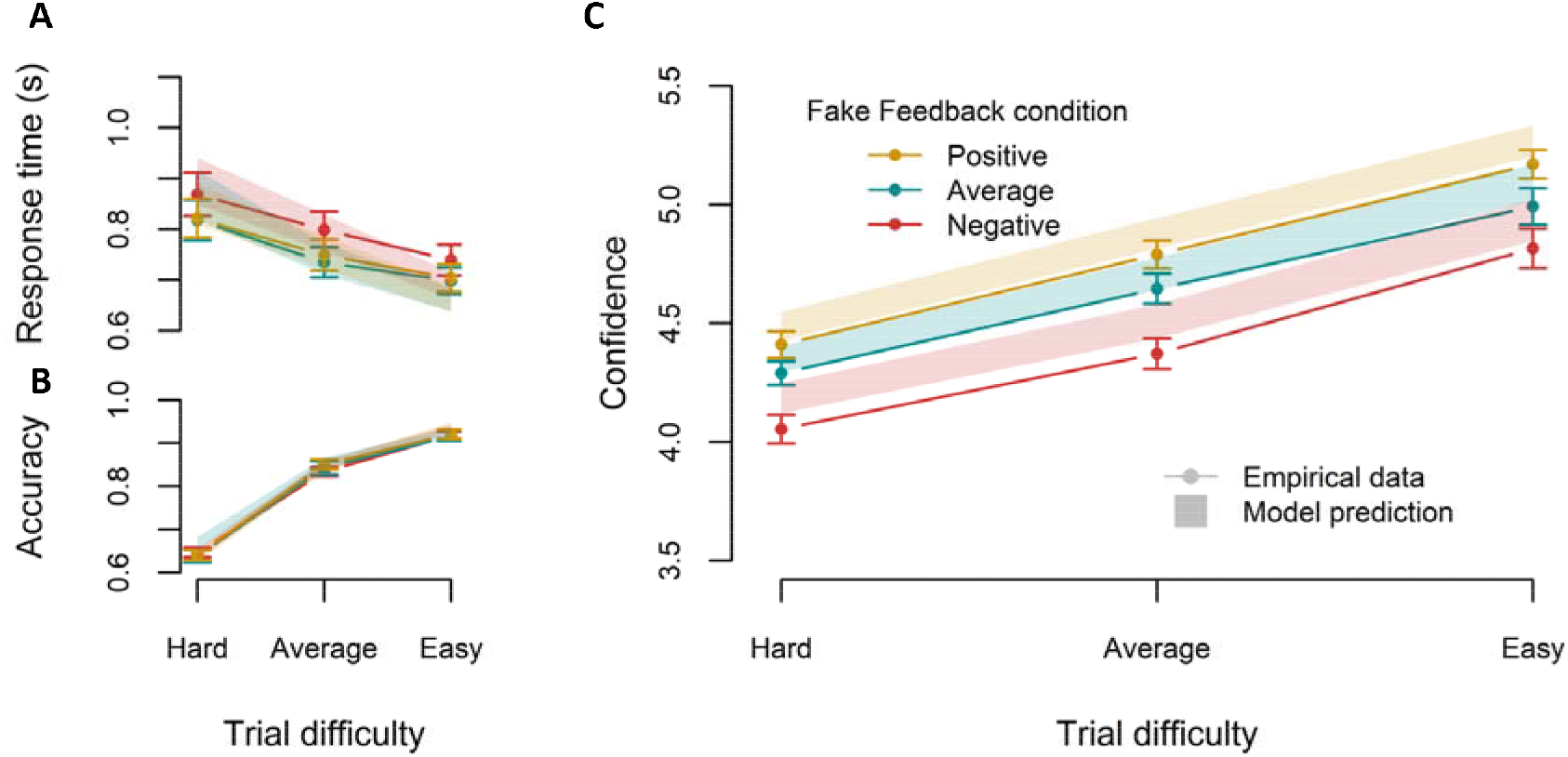
A key prediction of the LDC model is that confidence can vary independent from task performance. **A.** In Experiment 1, providing participants with fake feedback telling them their performance was better, equal or worse than a reference group indeed left RT unaffected. **B**. Same with accuracy. C. On the other hand, fake feedback selectively influenced the reported level of confidence on correct trials. These results were closely captured by fitting the LDC model to these data. *Note: Solid lines represent empirical data. Shades and error bars represent standard error of the mean for predictions of the LDC model and empirical data, respectively*.

We next investigated the estimated parameters of the model. Given that feedback selectively influenced confidence ratings, we expected a significant change in the confidence-specific parameters (i.e., α or β), but no variation in the DDM parameters (non-decision time, drift rate, decision threshold). Indeed, feedback had an influence on estimated α (F(2,382) = 6.56, p = .0016) and β (F(2,382) = 8.32, p < .001). Tukey’s test for multiple comparisons found that estimated α was lower in the negative condition than in the other two conditions (negative vs average: p = .01; negative vs positive: p = .002), whereas there was no difference in α between the average and positive conditions (p = .88). In a similar vein, β was higher in the positive condition compared to the other two (positive vs average: p = .004; positive vs negative: p < .001), whereas there was no difference between the negative and average conditions (p = .83). Finally, as expected estimated DDM parameters did not vary with feedback condition (drift rate: F(2,48) = .18, p = .84; non-decision time: F(2,382) = 0.99, p = .37) except for a minor effect on decision threshold (F(2,382) = 3.30, p = .038). Post-hoc tests for the decision threshold revealed a slightly higher threshold in the negative condition compared to the positive condition and no difference with the other contrasts (negative - average: p = .14; negative - positive: p = .04; average - positive: p = .85).

#### Experiment 2: Dissociating parameter-specific effects on confidence ratings

Our final aim was to demonstrate that humans are sensitive to the specific parameterization of decision confidence proposed by the LDC framework. If confidence is computed using a low-dimensional solution, it should be possible to independently manipulate its parameters. Therefore, in a new set of two experiments, we aimed to induce selective changes in each parameter (reliability (α) or bias (β)) of the model.

The general design of both experiments was similar to Experiment 1: we manipulated the feedback during a training phase and investigated the impact of that manipulation on confidence ratings reported in a subsequent testing phase. Rather than presenting fake feedback every 24 trials, we adopted a novel approach where feedback during the training phase was presented after each trial in the form of a continuous value (Figure 2). Participants were told that this value reflected the probability that their response was correct (e.g., .8 vs .4 indicating that there was a high vs low probability that they just made a correct choice). Unknown to participants the exact feedback value was generated by LDC behind the curtain (see Methods for full details). Both experiments comprise a baseline condition (α = 18; β = 0) in which the feedback presented to participants reflected the model-approximated probability of a choice being correct. In Experiment 2A, the value of α that was used to generate the feedback was selectively manipulated between conditions. In addition to the baseline condition there was a minus condition where α was decreased (α = 9), and a plus condition where α was increased (α = 36). In Experiment 2B, the same procedure was used except that now the value of β was selectively manipulated between conditions (β = −1 in the minus condition and β = 1 in the plus condition).

##### A dissociable effect of manipulated feedback on confidence according to the parameter manipulated

As previously described, the reliability parameter α reflects how strongly individuals weigh evidence in their computation of confidence. Given that accuracy is closely related to the amount of available evidence, correct trials tend to have considerable supporting evidence when reporting confidence, whereas error trials usually have little to no supporting evidence. Given that α weighs evidence, a decrease (in the α-minus condition) or an increase (in the α-plus condition) of α is therefore expected to differently impact confidence for correct trials (strong influence) than for error trials (little to no influence). In contrast, the parameter β reflects a stimulus-independent confidence bias, so providing participants with β-manipulated feedback is expected to lead to changes in confidence irrespective of choice accuracy. The reasoning for this prediction is that β is not concerned with the evidence provided by the stimulus (nor by the response), as it simply adds (in the β-plus condition) or subtracts (in the β-minus condition) a constant to the (logit of the) confidence judgment regardless of what happens during the trial.

These intuitions are further illustrated in Figure 4A and 4B, which show the actual (i.e., manipulated) feedback that was presented to participants during the training phase of our experiments. Confirming the above intuition, there was an interaction in Experiment 2A between accuracy and the value of α (F(2,12623) = 76.73, *p* <.001): feedback was more positive when generated by a higher α when considering correct trials only (F(2,9911) = 723.77, *p* < .001), but did not change when considering error trials only, F(2,44) = 1.41, *p* = .25. For Experiment 2B, although there was a significant interaction between β value and accuracy (F(2,10589) = 43.11, p <.001), feedback was more positive when generated by a higher β both when taking corrects (F(2,8420) = 2260.84, *p* < .001) and errors (F(2,35) = 183.56, *p* < .001) separately.

**Figure 4.**
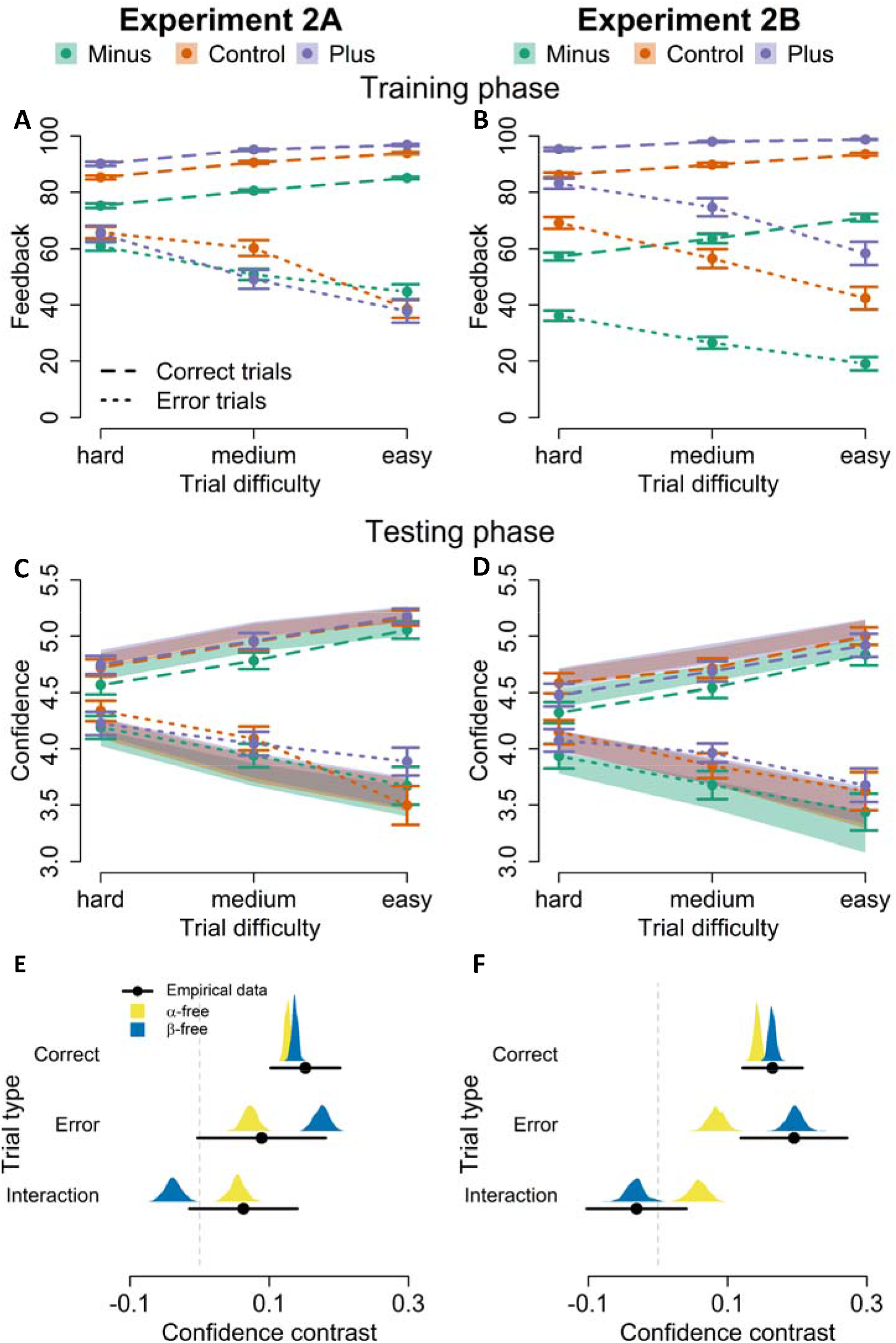
A key prediction of the LDC model is that participants should be sensitive to the specific parametrization of confidence proposed by the model. To test this, Experiment 2 provided participants with probabilistic feedback generated by the LDC model. Critically, LDC based feedback was generated using different levels of α or different levels of β. A. Changing α influences the confidence for correct trials but not for errors. B. Changing β influences feedback for both corrects and errors. The pattern that we saw in the feedback (which effectively are our predictions) was also seen in the behavioral data. C. α-manipulated feedback influenced confidence reports for correct but not error trials. D. β-manipulated feedback influenced confidence reports on both correct and error trials. E. Fitting the LDC model to the empirical data of Experiment 2 revealed that data in the α-manipulated feedback was best explained by a model in which α was allowed to vary. F. Data from the β-manipulated feedback was best explained by a model in which β was allowed to vary. To visualize this, we computed confidence contrasts for the empirical data (black lines), as well as for the α-free (yellow distribution) and β-free (blue distribution) model fit, separately for corrects and errors. “Interaction” refers to the difference between the confidence contrast in corrects and errors. *Note: black dots correspond to the average empirical contrast, distributions correspond to the bootstrapped mean predicted confidence contrasts. Error bars and shaded areas represent empirical and model-simulated SEM, respectively*.

##### Behavioral results

We now turn to the effects of the feedback manipulations on the testing phase data. The results concerning task performance were as expected: we found no effect of feedback condition on performance (RT and accuracy) in the testing phase of Experiment 2A (RT: F(2,40) = .15, p = .86; Accuracy: *X*^2^(2) = 3.54, p = .17) and Experiment 2B (RT: F(2,32) = .24, p = .79; Accuracy: *X*^2^(2) = 1.09, p = .58). There was, however, the expected effect of trial difficulty on performance both in Experiment 2A (accuracy: *X*^2^(2) = 1023.00, *p* < .001; RT: *F*(2,25619) = 164.34, *p* < .001) and Experiment 2B (accuracy: *X*^2^(2) = 767.30, *p* < .001; RT: *F*(2,21189) = 170.16, *p* < .001), with lower accuracy and higher RT when trial difficulty was higher (all post-hoc comparisons: ps < .02). There was no interaction between feedback condition and trial difficulty on RT and accuracy in either Experiment 2A or 2B (all ps > .05).

After demonstrating that the feedback did not influence task performance itself, we next turn towards confidence ratings. In line with the feedback presented during the training phase (Figure 4A and 4B), the data of the testing phase revealed that α-manipulated feedback in Experiment 2A had an effect on confidence ratings within correct trials (F(2,39) = 4.86, p = .01), but not within error trials (F(2,45) = .87, p = .43; Figure 4C). Note that this finding should be interpreted with caution, since the interaction between accuracy and feedback was not significant (F(2,44) = .62, p = .54). Turning to Experiment 2B, in line with the predictions there was an effect of feedback condition on confidence ratings in both correct trials (F(2,33) = 8.86, *p* < .001) and in error trials (F(2,36) = 4.28, p = .02; Figure 4D). Here again, no interaction between accuracy and feedback condition was found (F(2,35) = .29, p = .75). Lastly, trial difficulty had an effect on confidence ratings in both Experiment 2A (F(2,25633) = 75.21, p < .001) and 2B (F(2,253) = 33.49, p < .001). We found no interaction between trial difficulty and feedback condition (Experiment 2A: F(4,25625) = 2.37, p = .05; Experiment 2B: F(4,20385) = 1.70, p = .15). Overall, these results corroborate the predicted pattern and show a clearly dissociable effect of feedback on confidence ratings according to the parameter manipulated in the feedback generation.

##### LDC model fits

We next performed model comparison to explore whether the different patterns of confidence ratings observed in Experiment 2A and 2B would be best explained by a change in the targeted parameter (i.e. a change in α in Experiment 2A and a change in β in Experiment 2B). Two candidate LDC models were fit to the accuracy, RT and confidence data of both experiments. Each model differed in whether α or β was fixed between feedback conditions: in the α-free model, only α was allowed to vary between feedback conditions, whereas in the β-free model, only β was allowed to vary between feedback conditions. As recommended in Palminteri et al. (2017), we investigated how well simulations from the best-fitting parameters from both the α-free and the β-free models were able to reproduce the observed behavioral effects. Specifically, we defined a confidence contrast that captured the qualitative signatures seen in the feedback presented. Since the difference in feedback between the baseline and the plus condition was negligible relative to how both conditions differed from the minus condition in both experiments, we computed our confidence contrast as average confidence in the minus condition subtracted from average confidence in the baseline and the plus condition. Figure 4E-F show the empirical confidence contrast as well as the distribution of the mean predicted confidence contrast for both the α-free and the β-free model obtained via bootstrapping. In Experiment 2A, the confidence contrasts predicted by both the α-free and the β-free model was highly similar for correct trials, and both matched well to the empirical data. However, while the α-free model closely captured the confidence contrast in errors and hence the interaction, the β-free model overestimated the effect in errors, which led it to underestimate the interaction. Similarly, in Experiment 2B, both models accurately captured the empirical confidence contrast in correct trials. Additionally, the β-free model nicely reproduced both the empirical confidence contrast in error trials and the interaction, whereas the α-free model clearly underestimated the confidence contrast in error trials, which led to predicting an interaction that was not present in the empirical data.

To further confirm that the α-free (resp. β-free) model is the most likely to explain the results of Experiment 2A (resp. 2B), we additionally quantified the goodness-of-fit of each model using Bayesian information criterion (BIC). Two additional candidate models were included in that analysis: a null model where neither α nor β varied between feedback conditions and a full model where both α and β were free to vary between conditions. Table 1 reports the difference in BIC of each candidate model compared to the best model, separately for both experiments. A first conclusion that can be drawn, is that both the α-free and β-free model outperformed the null model (i.e., providing strong evidence for a change in the parameters) as well as the full model (i.e., providing strong evidence for a *selective* change in the parameters). Second, as expected the α-free model showed the lowest BIC for the data of Experiment 2A. Surprisingly though, the α-free model also slightly outperformed the β-free model in Experiment 2B. Overall, the difference in BIC between the α-free and the β-free models appears marginal compared to how strongly they each outperformed the null and full models. Additionally, the difference in BIC between the α-free and the β-free models was bigger in Experiment 2A (Δ_*BIC*_ = 4.15), where the α-free model was expected to be the best performing model, compared to the difference observed in Experiment 2B (Δ_*BIC*_ = 2.54). Applying categorical cutoffs to describe the magnitude of the evidence in favor of the α-free model in both experiments, such as the rule of thumb proposed by Burnham & Anderson (2004), leads to conclude that the α-free model has considerably more support than the β-free model in Experiment 2A, but only weak support in Experiment 2B. Taken together, these results suggest that theoretically motivated confidence manipulations can lead to specific and theoretically predicted changes in confidence.

**Table 1.**
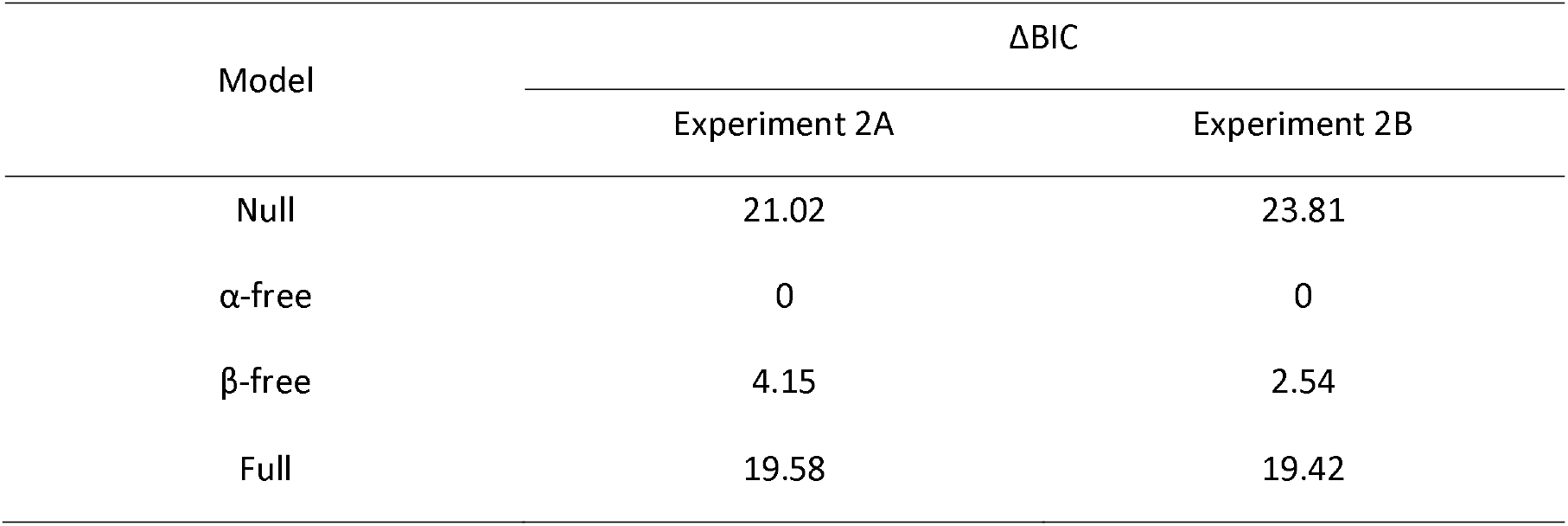
Model comparison expressed in distance in BIC from the best-fitting model.

## Discussion

How to incorporate the sense of confidence in models of decision-making has been the focus of much recent work. An influential framework is based on the Bayesian interpretation of confidence (3, 32–34), namely that confidence reflects the probability of being correct given both accumulated evidence and elapsed time (14, 15, 17). In order to accurately compute this probability, it is necessary to know how to compute confidence based on the available data (evidence and time). Currently, a computationally plausible account describing how individuals learn this mapping is lacking. In the current work, we introduced the LDC model, which provides a tractable and flexible account of decision confidence. Using simulations, we first showed that LDC provides a highly reliable approximation of the true probability correct. Fitting this model to empirical data revealed that LDC accounts very well for human confidence ratings. Critically, using a novel feedback manipulation, we validated two key predictions from the model, namely that 1) changes in confidence can be independent of performance and 2) independently manipulating the reliability (α) and bias (β) parameters elicit clearly dissociable and identifiable effects on confidence.

### Introducing tractability and flexibility to decision confidence modelling

The LDC model belongs to the family of DDM-based models of decision confidence. Here, confidence is conceptualized as a (Bayesian) readout of the probability of a correct choice given evidence, time and choice. Existing models following that approach have been successful in explaining a wealth of data, including the link between confidence and RT (14, 17), and deviations from accuracy through the contribution of priors (18, 19). Estimating the probability correct based on the available data, however, is computationally intractable. The LDC model therefore proposes to approximate the Bayesian readout with a logistic function, offering a tractable approach of how humans compute confidence.

To increase flexibility and account for deviations from optimality, the LDC model relies on two free parameters, which control the reliability of evidence (α) and general biases (β) in the computation of confidence. A different class of confidence models that can account for biases and deviations between confidence and accuracy is based on Signal Detection Theory (SDT) framework (35–40). These models typically either assume the existence of metacognitive noise (37–39), and/or consider that confidence is not entirely derived from the same signal as the primary decision (35–38, 40). A recent study comparing the different SDT models of confidence on simple perceptual tasks showed that confidence is simply computed as a noisy readout of the evidence used for the primary decision (41). Although the LDC model is grounded within the DDM tradition which conceptualizes confidence as the Bayesian probability correct, it does not critically hinge upon the specifics of the DDM. It would be straightforward to construct a simplified version of the LDC model which ignores the element of time. This would allow to directly compare the LDC approach to recent SDT models of confidence. Crucially, with its parameters, our model can flexibly account the different types of idiosyncrasies, biases and deviation from the optimal Bayesian readout (21–24), which are all merged into a single metacognitive noise parameter in most SDT frameworks.

### Confidence can vary independently from task performance

In both Experiments, we observed that decision confidence was influenced by the feedback manipulation, whereas objective performance was not. This rules out an interpretation whereby the feedback influenced task performance and changes in confidence simply reflect this change in performance. Indeed, some previous work has shown that changes in confidence can be explained by subtle differences in RT (14, 42). This was not the case in the current experiments. As such, it is unlikely that Bayesian read-out models can account for the effects observed in the current work, as they do not allow for confidence-specific parameters (14–16; for a counterexample see 17). In contrast, LDC nicely captured the effect of feedback on confidence in the absence of changes in objective performance, thus attesting to the flexible nature of the LDC model. Previous studies have unraveled several other factors that influence the reported level of decision confidence, while leaving task performance unaffected, for example emotional states (27, 43), working memory content (29) and age (44, 45). More broadly, dissociations between performance and metacognition have long been reported in cases such as blindsight (10, 46), where individuals with lesions in primary visual cortex show above chance level performance at visual tasks despite reporting no awareness of the stimuli. At the opposite end of the spectrum, change blindness (i.e. failure to detect major differences between two images while they flicker off and on) is a typical example of a metacognitive error where individuals believe they would be able to detect such major changes, despite being unable to do so (11). These examples highlight how ubiquitous dissociations between performance and metacognition are. By incorporating free parameters controlling for evidence reliability and bias into the computation of confidence, the LDC model is in principle flexible enough to account for all these dissociations reported in the literature.

### Humans can independently tune evidence reliability and bias in confidence

In Experiment 2A and 2B, we aimed to selectively manipulate confidence ratings according to each parameter of the LDC. By providing model-generated feedback from different α’s in Experiment 2A and different β’s in Experiment 2B, we revealed clearly distinct patterns of confidence ratings according to the parameter manipulated. Moreover, the empirically observed patterns were best captured by models where the manipulated parameter was set as a free parameter (e.g. α-free model when feedback was α-manipulated). These results imply that individuals can change their computation of confidence consistently with our parameterization of confidence, providing strong validating evidence in favor of LDC. This observation raises the intriguing possibility that individuals might exert control over the parameters governing the computation of confidence in a way that maximizes utility. Intuitively, computing confidence in such a way that it closely matches the Bayesian readout seems like the rational strategy to optimize utility, as it would allow to optimize behavior based on the best possible internal evaluation of that behavior (5, 7, 9). In some contexts, however, other factors than informativeness play a role in the utility of confidence. When competing for shared limited resources, expressing overconfidence plays a key role in convincing other agents not to compete for the resource (i.e. “bluffing”; 47, 48). Errors caused by overconfidence, though, bear a high cost in such strategy. In such a context, the optimal way to compute confidence seems to be an increase in the evidence reliability estimate (α), which will lead to higher confidence for scenarios with much evidence (i.e., overconfidence when you are likely to win the competition) but lower confidence for scenarios with little evidence (i.e., when you are likely to lose the competition). Increasing β in this scenario is likely suboptimal because this produces overall high confidence, also for scenarios with little evidence. The opposite scenario might be true in a social decision-making context. If confidence is used to assert influence rather than to convey accuracy (49), the optimal strategy might be an overall increase in β, resulting in general overconfidence (i.e. irrespective of accuracy) to push forward one’s choice. These examples show that what is traditionally treated as deviations from the optimal Bayesian readout can sometimes be considered as optimal through the lenses of utility maximization.

### Beyond dichotomies with model-informed feedback

In contrast with the binary “correct/error” feedback typically provided in lab experiments, feedback received in daily life is not always clear-cut. Individuals must often make sense of noisy and probabilistic feedback cues (e.g. how should a street-artist interpret a subtle nod from a spectator?). Continuous feedback has been used in the past to communicate performance relative to other (hypothetical) participants (19, 50) or to give average accuracy over several past trials (51, 52). However, in the current work we designed a novel feedback manipulation which provides continuous feedback about choice accuracy on a trial-by-trial basis. It is important to note that our instructions simply stated that feedback would reflect the probability of being correct on a single trial, without much more explanation as to how this proportion was calculated. A skeptical participant could reasonably doubt the trustworthiness of the feedback, since it might seem unlikely that we provide an “accurate” probability of being correct on a single trial basis (e.g. is a feedback of 80% vs 70% really informative, or are the values pure noise added by the experimenter). Despite these potential obstacles, our feedback manipulation did produce the confidence patterns we predicted, hence validating our model-generated feedback approach. This nuanced way of providing feedback goes beyond the mere distinction between dichotomous valid versus invalid feedback (53), and offers a promising framework to control the level of ambiguity and informativeness of trial-by-trial feedback, allowing to study in a more fine-grained manner how individuals process and are impacted by more realistic, ambiguous feedback (54, 55).

### Interpreting the LDC parameters

An appealing property of computational models is that their parameters often have clear interpretations, and can be selectively manipulated (13, 56), although it is subject of recent debate (57). Similarly, in LDC, evidence and time are mapped onto confidence by means of a reliability parameter, α, and a confidence bias parameter, β. Our reliability parameter, α, can be interpreted as an individual’s estimate of the precision of evidence. This interpretation is similar to the recently proposed concept of “meta-uncertainty”, which is described as “the subject’s uncertainty about the uncertainty of the variable that informs their decision” (58). In both the LDC model and Boundy-Singer et al.’s CASANDRE model, one’s estimate of evidence reliability weighs how evidence is used to compute confidence. Note that an important difference is that in CASANDRE the estimate is assumed to be correct on average (i.e. individuals are assumed to have an uncertain, but on average correct estimate of evidence reliability), whereas one of the key points of the LDC model is that participants can have incorrect values of α.

The second parameter of LDC, β, globally increases or decreases confidence. It straightforwardly relates to the metacognitive bias described in other models of confidence (59). In light of this interpretation of α and β, one can further interpret specific patterns in the data. For example, in Experiment 1, we observed a change in α in response to negative feedback (with a significantly lower estimated α compared to the other two conditions), indicating that participants judged evidence as less reliable after receiving negative feedback. On the contrary, we observed a change in β after positive feedback (with a significantly higher estimated β compared to the other two conditions), suggesting a general overconfidence bias after receiving positive feedback. This dissociation suggests that despite similar effects at the behavioral level, the LDC model allows to further tease apart the origins of confidence biases e.g. in response to positive vs negative feedback.

Finally, we note that in the current parameterization of confidence, identical to the Bayesian readout, confidence always depends on 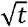. However, the influence of time on confidence might vary according to the task or individual. To account for such hypothetical sources of variability, one could expand the LDC model by further parameterizing the influence of time with a third parameter, γ, and replace 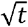 in Eq. (3) with *t*^γ^. The model then has an accurate calibration of how time influences confidence when γ = 0.5, and overweighs (resp. underweighs) time in the computation of confidence when γ > 0.5 (resp. γ < 0.5). By doing so, future work might investigate whether variability in the relation between confidence and decision time can be captured by the extended LDC model.

## Conclusion

We introduced the LDC model, a new model of decision confidence that offers a tractable and flexible approximation of confidence as the Bayesian probability of making the correct decision. The model provides a low-dimensional parametrization of decision confidence which allows efficient estimation of confidence, while at the same time accounting for idiosyncrasies and different kinds of confidence biases. This parameterization of confidence was validated in two experiments showing a distinct pattern of confidence ratings after specifically manipulating the mapping according to each parameter of the model.

## Methods

### Experiment 1

#### Participants

Fifty participants (eight men, one third gender, age: M = 19, SD = 4.9, range 17–52) took part in Experiment 1 (two excluded due to chance level performance). All participants participated in return for course credit and read and signed a written informed consent at the start of the experiment. All procedures were approved by the KU Leuven ethics committee. Detailed methods and analyses for Experiment 1 have already been reported in Van Marcke et al. (2022). We briefly report the general procedure here.

#### Procedure

Participants completed three decision-making tasks: a dot color task, a dot number task and a letter discrimination task. Each task started with 120 training trials. Feedback during training was presented at the end of blocks of 24 trials. Unknown to participants, feedback was predetermined to be either good, average or bad for a specific task, and feedback scores were randomly sampled according to the feedback condition. Each participant received good feedback on one task, average feedback on another task, and bad feedback on a third task (order and mapping with tasks counterbalanced between participants). After the training phase of a task, participants performed 216 test trials where feedback was no longer provided. Instead, confidence ratings were queried at the end of each trial. For each task, there were three levels of stimulus difficulty (easy, average, or hard).

##### Dot color task

On each trial, participants decided whether a box contained more (static) blue or red dots. The total number of dots was always 80, with differing proportions of red or blue dots depending on the difficulty condition. The position of dots was randomly generated on each trial.

##### Dot number task

On each trial, two boxes were presented, one of which contained 50 dots and the other more or less than 50 dots. Participants decided which of the two fields contained the largest number of dots. The exact number of dots in the variable field differed depending on the difficulty condition. The position of dots was randomly generated on each trial.

##### Letter discrimination task

On each trial, participants decided whether a field contained more X’s or O’s. The total number of X’s and O’s was always 80, with differing proportions of X’s or O’s depending on the difficulty condition. The position of the letters was randomly generated on each trial.

### Experiment 2

#### Participants

Forty-three participants (8 male, age: M=18.49, SD=1.03, range 16-22) took part in Experiment 2A. Forty-two participants (9 male, age: M=18.83, SD = 2.05, range 17-29) took part in Experiment 2B. Due to chance performance on at least one of the tasks, we removed 3 participants from Experiment 2A and 3 participants from Experiment 2B. Six additional participants were removed from Experiment 2B due to (almost) no variability in their confidence reports (i.e. used the same report on more than 90% of the trials). All participants took part in return for course credit and signed informed consent at the start of the experiment. All procedures were approved by the local ethics committee.

#### Stimuli and apparatus

All experiments were conducted on 22-inch DELL monitors with a 60 Hz refresh rate, using PsychoPy3 (Peirce et al., 2019). All stimuli were presented on a black background centered on the middle of the screen (radius 2.49° visual arc). Stimuli for the dot number task were presented in two equally sized boxes (height 20°, width 18°) at an equal distance from the center of the screen. Stimuli for all other tasks were presented in one box (height 22°, width 22°).

#### Procedure

In both experiments, participants completed three decision-making tasks: a dot color task, a shape discrimination task and a letter discrimination task. Each task started with 108 training trials. After each choice, participants rated their confidence level and then received (continuous) feedback about their performance. After the training phase of a task, a test phase of 216 trials followed which was identical to the training phase, except that feedback was omitted. Every trial was assigned one of three possible difficulty levels. The difficulty levels were matched between the three tasks based on a pilot staircase session. For all tasks, a trial started with a fixation cross that was presented for 500 ms, after which the stimulus appeared for 200 ms or until a response was given. Participants indicated their choice using the C or N key using the thumbs of both hands. There was no time limit for responding, although participants were instructed to respond as fast and accurately as possible. After each choice, participants rated their confidence on a 6-point scale, labeled from left to right: ‘sure error’, ‘probably error’, ‘guess error’, ‘guess correct’, ‘probably correct’, and ‘sure correct’ (reversed order for half the participants). Confidence was indicated using the 1, 2, 3, 8, 9 and 0 keys at the top of the keyboard with the ring, middle and index fingers of both hands. There was no time limit for indicating confidence. During the training phase only, a trial ended with a visual presentation of feedback. An empty horizontal rectangle was filled in white starting from the left end of the rectangle (reversed order for half the participants, matched to the confidence counterbalancing). The proportion filled corresponded to the probability that the response was correct (e.g. halfway filled if feedback is 50%). Ticks at the 0, 25, 50, 75 and 100 percent marks were respectively labeled ‘sure error’, ‘probably error’, ‘random chance’, ‘probably correct’ and ‘sure correct’.

On each trial, participants decided whether a box contained more elements from one out of two categories. In the letter discrimination task, elements were A’s or B’s, in the dot color task, blue or red dots and in the shape discrimination task, squares and circles. The total number of elements in a box was always 80, with the exact proportion of each element depending on the difficulty condition. The position of the elements was randomly generated on each trial.

#### Model-generated feedback

Instead of binary feedback (correct/error), feedback during the training phase after each trial was provided in the form of a continuous value. Participants were told that this probability reflected the probability that their response was correct. In reality, the feedback was generated by our model of confidence. To do so, we estimated the single-trial evidence accumulation process online (i.e., during the experiment). To do so, we assumed that performance was equivalent to the average performance observed in piloting sessions. In other words, we assumed that the current decision threshold and drift rate were equal to the average decision threshold and drift rate from piloting sessions. At the moment a decision was made, the evidence accumulation process just reached the decision threshold. We thus inferred that the amount of accumulated evidence at the time of decision was equal to the average decision threshold estimated from the pilot sessions. Then, to estimate the total amount of accumulated evidence at the time of the confidence report, we added the post-decisional evidence estimated by running a random-walk for a duration fixed to the observed confidence RT and with a drift rate set to the average drift rate estimated from the pilot sessions (the sign of which varied whether the response was correct or not). Feedback was thus equal to model confidence computed according to a fixed (α, β) pairing (the value of which depended on the condition and experiment one is in) from that total evidence and the total time (decision RT + confidence RT).

#### Feedback conditions

In a baseline condition, the feedback presented to participants reflected the actual model-generated probability of a choice being correct. To get the value of α and β that best approximate the true probability of a choice being correct, we estimated both parameters based on the heatmap generated by the drift rates observed in the pilot sessions. In the baseline condition, α was thus set to 18 and β to 0. In Experiment 2A, for one task feedback was computed using a lower value of α (namely 9), and for another task feedback was computed using a higher value of α (namely 36; termed “α-plus” condition). The association between the manipulation of α and the task was counterbalanced across participants. In Experiment 2B, feedback was provided according to the baseline condition in one task, using a lower value of β in another task (−1), and using a higher value of β in another task (1).

#### Statistical analyses

All data were analyzed using mixed effects models. We started from models including the fixed factors and their interaction(s), as well as a random intercept for each participant. These models were then extended by adding random slopes, only when this significantly improved model fit. Confidence ratings and RT were analyzed with linear mixed effects models, for which we report *F* statistics and the degrees of freedom as estimated by Satterthwaite’s approximation. Accuracy was analyzed using a generalized linear mixed model, for which we report *X^2^* statistics. All model fit analyses were done using the lmerTest R package (60).

#### Bounded evidence accumulation

We modeled choice and RT data using the drift diffusion model (DDM), a popular variant of the wider class of accumulation-to-bound models. In the DDM, noisy evidence (representing the difference between the evidence for both options) is accumulated, the strength of which is controlled by a drift rate *v*, until one of two decision thresholds *a* or -*a* is reached. Non-decision components are captured by a non-decision time parameter *ter*. To simulate data from the model, random walks were used as a discrete approximation of the continuous diffusion process of the drift diffusion model. Each simulated random walk process started at *z*a* (here, *z* was an unbiased starting point fixed to 0). At each time step *τ*, accumulated evidence changed by Δ with Δ given in Eq. (3):

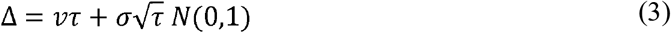

Within-trial variability is given by *σ*. In all simulations, *τ* was set to 1 ms, and *σ* was fixed to .1.

#### Model fitting

Model predictions were obtained from the random-walk simulation described above. Evidence continued to accumulate after threshold crossing for a duration that was sampled from the confidence RT distribution of the trials being fitted. Note that this sampling was done without replacement, ensuring that the simulated confidence RT distribution exactly matched the empirically observed confidence RT distribution. The number of trials being simulated was equal to 20 times the number of empirical trials being fitted to ensure that every trial of the empirical confidence RT distribution is being simulated an equal amount of time. Given that the model-generated confidence comes on a continuous scale from 0 to 1, we binned the model output into 6 equally-spaced bins.

Accuracy and RT data of each task and participant was estimated using 5 DDM parameters: non-decision time, decision threshold and three drift rate parameters (one for each trial difficulty level). Additionally, α and β were fitted to the confidence judgments, separately for each feedback condition. We implemented quantile optimization, and computed the proportion of trials falling within each of six groups formed by quantiles .1, .3, .5, .7 and .9 of RT, separately for corrects and errors. Similarly with confidence ratings, we computed the proportion of trials resulting at each of the 6 levels of confidence judgment separately for corrects and errors. The resulting objective function consisted in minimizing the sum of squared errors described in Eq (4):

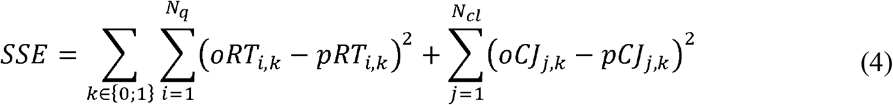

with *N_q_* = *N_cl_* = 6 the number of RT groups/possible confidence value, *oRT_i,k_* and *pRT_i,k_* respectively the proportions of observed and predicted trials falling within quantile i of RT, separately for corrects (*k* = 1) and errors (*k* = 0), and *oCJ_i,k_* and *pCJ_i,k_* reflecting their counterpart for confidence. Models were fitted using a differential evolution algorithm (61), as implemented in the DEoptim R package (62). The population size was set to 10 times the number of parameters to estimate. The algorithm stopped once no improvement of the objective function was observed for the last 100 generations.

#### Model comparison

All candidate models for the model comparison were based on the same estimated DDM parameters fitted separately to accuracy and RT data (i.e. minimizing the first term of the SSE in Eq. (4)). Each candidate model was then fit to confidence ratings (i.e. minimizing the second term of the SSE in Eq. (4)). BIC values for model comparison were computed as follows (63):

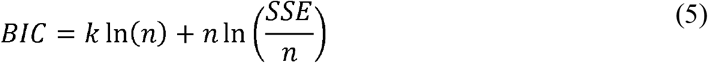

with *k* the number of free parameters and *n* the number of data points. BIC values for each model represented in Table 1 correspond to the mean BIC over participants. Bootstrapped 95% confidence intervals of confidence contrasts were obtained by simulating 500 datasets based on the fits of each participant and then computing the mean confidence contrasts of each repetition. The 95% confidence interval was computed as the .025 and .975 quantiles of the distribution formed by the bootstrapping.

## Code availability

All raw data and analysis code can be freely accessed at https://github.com/pledenmat/ldc_paper.

## Acknowledgements

The research was supported by a starting grant from the KU Leuven (STG/20/006), a Francqui Start-Up Grant (PXF-D8830) and a grant from the Research Foundation Flanders, Belgium (FWO-Vlaanderen) (G0B0521N).

## Notes

https://github.com/pledenmat/ldc_paper

### Competing Interest Statement

The authors have declared no competing interest.

